# Physiology and lipid accumulation capacity of different *Yarrowia lipolytica* and *Rhodosporidium toruloides* strains on glycerol

**DOI:** 10.1101/278523

**Authors:** Suellen Patricia Held Azambuja, Nemailla Bonturi, Everson Alves Miranda, Andreas Karoly Gombert

## Abstract

**Objective:** To compare physiological and process parameters, as well as lipid accumulation capacity, of six strains of *Yarrowia lipolytica* and two strains of *Rhodosporidium toruloides* in media containing glycerol as the main carbon and energy source.

**Results:** The strains *Y. lipolytica* IMUFRJ 50678, Po1g, W29 and CCT 5443 displayed very similar physiological parameters, with µ_max_, 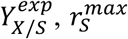 and 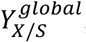 average values of 0.31 h^−1^, 0.53 (g DW/g S), 0.60 (g S/g DW.h) and 0.46 (g DW/g S), respectively. The two strains of *R. toruloides* presented physiological and process parameters with no significant difference, with average values of 0.084 h^−1^, 0.53 (g DW/g S), 0.17 (g S/g DW.h) and 0.44 (g DW/g S). Among all *Y. lipolytica* strains, *Y. lipolytica* CCT 5443 strain presented the highest *Y*_*Lip/S*_, 0.054 (g Lip/g S), and *P*_*Lip*_of 0.040 (g Lip/l.h). Among all investigated strains (*Y. lipolytica* and *R. toruloides*), the yeast *R. toruloides* CCT 7815 displayed the highest lipid accumulation capacity, with *Y*_*Lip/S*_equal to 0.11 (g Lip/g S) and *P*_*Lip*_equal to 0.10 (g Lip/l.h).

**Conclusion:** Among all strains investigated in our study, the yeast strain *R. toruloides* CCT 7815 presents the most promising characteristics for industrial single cell oil production.

**List of abbreviations:**

DW: dry weight (g/l)
Lip: lipid (g/l)
*P*_*Lip*_: lipid productivity (g Lip/l.h)
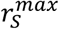: maximum specific substrate consumption rate (g S/g DW.h)
S: substrate (g/l)
*Y*_*Lip/S*_: lipid yield on substrate (g Lip/g S)
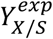: biomass yield on substrate during the exponential growth phase (g DW/g S)
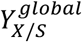: global biomass yield on substrate (g DW/g S)
µ_*max*_: maximum specific growth rate (h^−1^)

## Introduction

Microbial lipids, called single cell oil (SCO), has recently been gaining attention from industries due to its potential to be used for the production of cosmetics, pharmaceuticals, paints, other chemicals (Koutinas et al. 2014), and biodiesel (Papanikolaou and Aggelis 2011). Oleaginous yeasts are capable of accumulating at least 25 % of lipids in their dry biomass (Beopoulos et al. 2009). Among these yeasts, *Yarrowia lipolytica* and *Rhodosporidium toruloides* have been widely studied as SCO producers due to their ability to assimilate and convert into lipids a wide variety of substrates, including glycerol (Shi and Zhao 2017; Park et al 2018). Moreover, these yeasts can also be employed to obtain enzymes, organic acids, biosurfactants, among others compounds (in the case of *Y. lipolytica*) (Fickers et al. 2005), and carotenoids (in the case of *R. toruloides*) (Lee et al. 2014).

Most of the studies reported hitherto on SCO production using these two yeast species aimed at evaluating, in a systematic manner, their lipid accumulation capacity (Table 1), without looking at their basic physiology. Studies performed in defined media are rare or even absent. It is known that different strains, belonging to the same species, may present different properties. Here we present a systematic study of six strains of *Y. lipolytica* and two *R. toruloides* strains, in order to evaluate the following: 1) their physiological properties and potential to be used as SCO producers; 2) whether a defined medium is suitable for the growth of all strains; 3) how a hemicellulosic hydrolysate-evolved strain of *R. toruloides* performs in comparison with its parental strain. Glycerol was chosen as the main carbon and energy source in all cases, because it is an abundant by-product generated in the biodiesel industry (Lee et al. 2014).

**Table 1.**
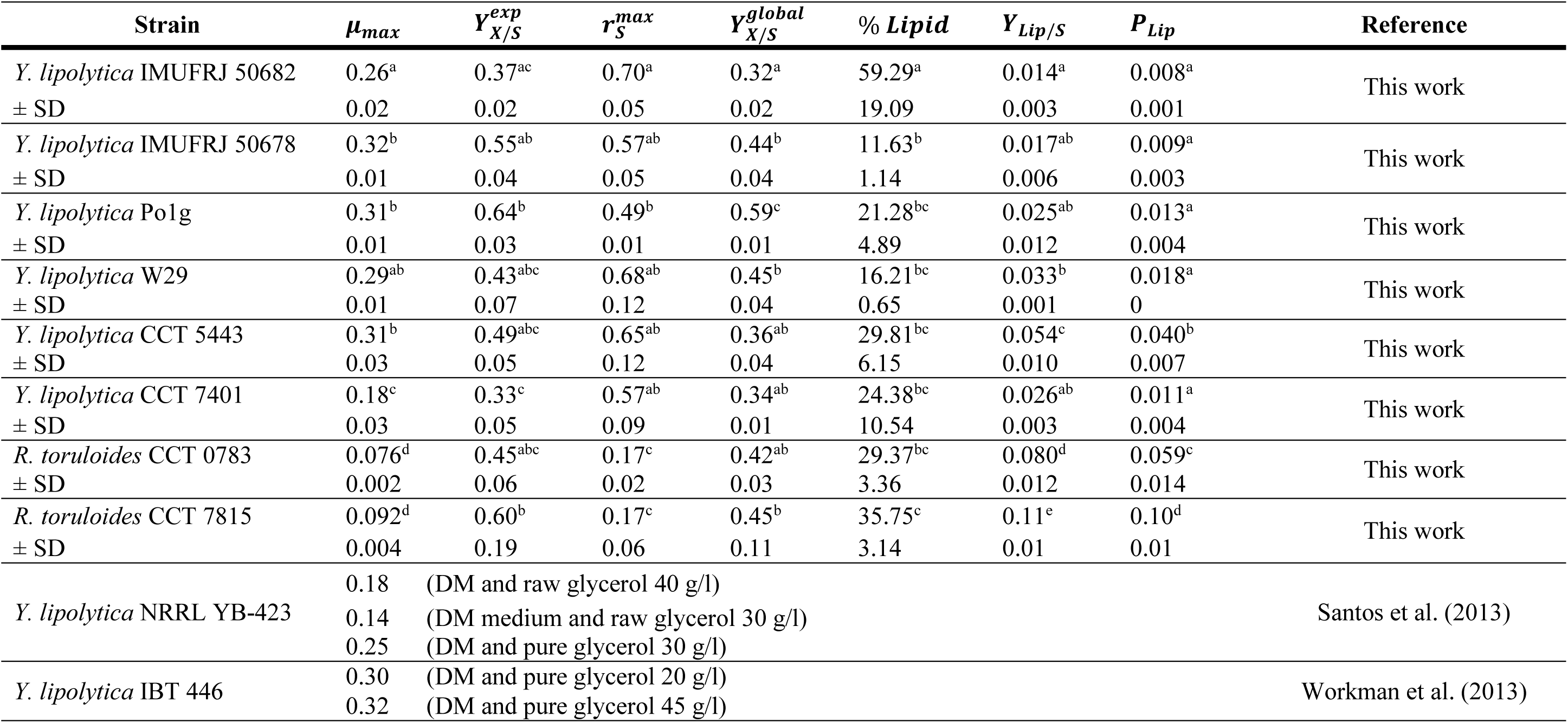
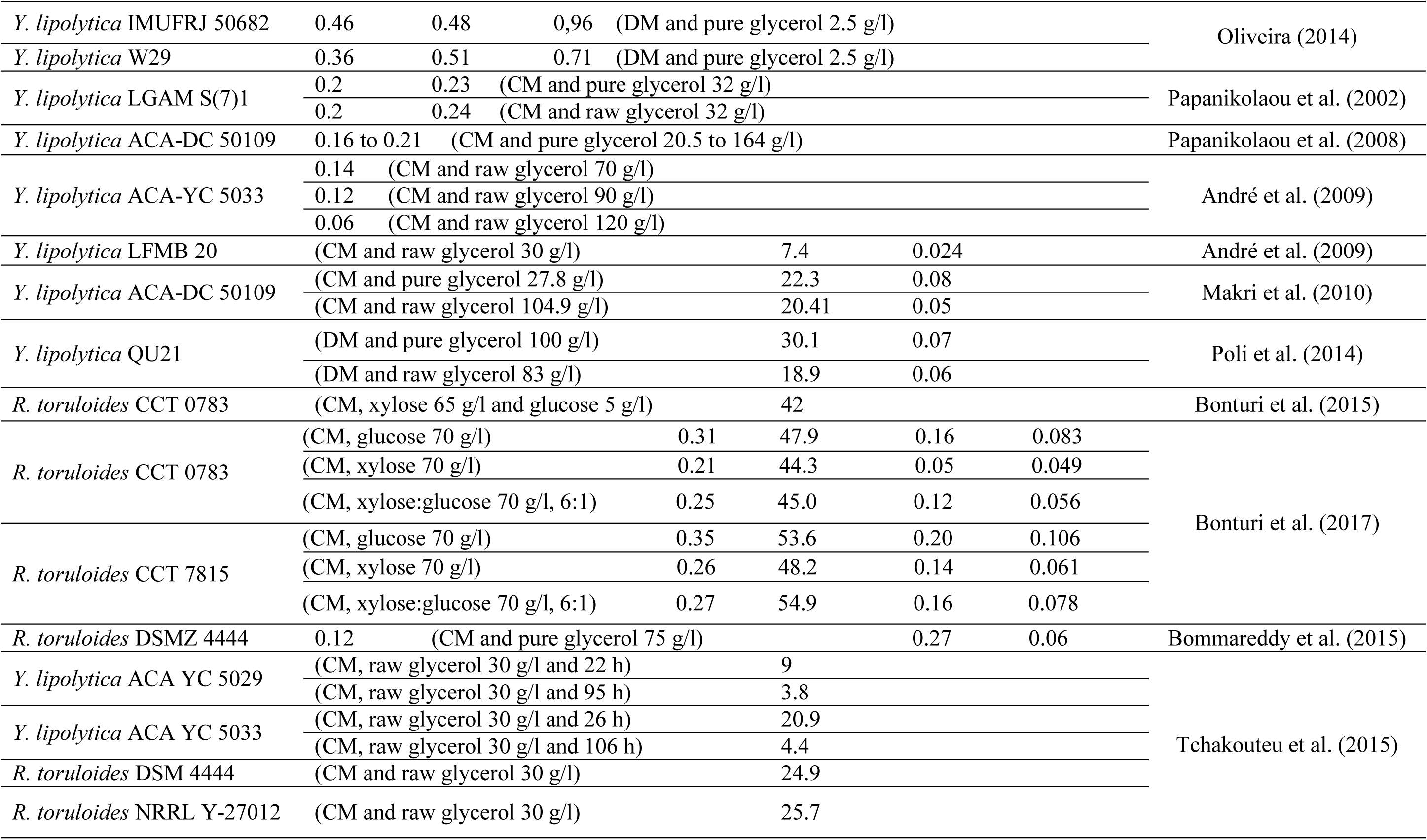
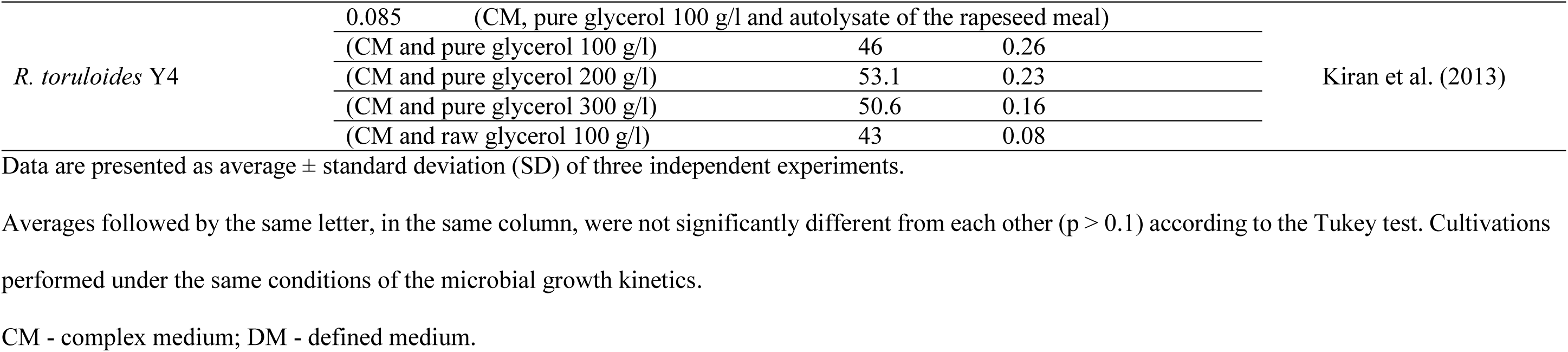
Maximum specific growth rate (µ_*max*_, h^−1^), biomass yield on substrate during the exponential phase (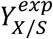, g DW/g S), maximum specific substrate consumption rate (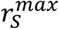, g S/g DW.h), global biomass yield on substrate (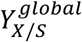, g DW/g S), lipid yield on substrate (*Y*_*Lip/S*_, g Lip/g S) and lipid productivity (*P*_*Lip*_, g Lip/l.h) during batch cultivations of *Y. lipolytica* and *R. toruloides* using glycerol as a sole carbon source (in this work) and different carbon and nitrogen sources (for other works)

## Materials and methods

Yeast strains and medium composition

The following *Y. lipolytica* strains were used: IMUFRJ 50682, IMUFRJ 50678, W29 and Po1g (Ura^−^ Leu^−^) (obtained from Prof. Maria Alice Zarur Coelho, Federal University of Rio de Janeiro, Rio de Janeiro, Brazil), CCT 5443 and CCT 7401 (Fundação André Tosello, Campinas, Brazil) (University of Campinas, Brazil). As for *R. toruloides*, a wild-type strain CCT 0783 (Fundação André Tosello, Campinas, Brazil) and its evolved derivative were used. The latter, named CCT 7815, was obtained after successive cultivations in increasing concentrations of sugarcane bagasse hydrolysate (5.3 g/l xylose, 0.4 g/l glucose, 1.4 g/l acetic acid, 0.2 g/l HMF, and 1.7 g/l furfural) (Bonturi et al. 2017).

The liquid defined medium MDBU used for determining growth kinetics consisted of (g/l): KH_2_PO_4_, 2.5; MgSO_4_7H_2_O, 0.5; and urea, 3.0. The MDBU medium was supplemented with 1 ml/l of thiamine solution (0.3 g/l) and 1 ml/l of trace element solution according to Barth and Kunkel (1979) for *Y. lipolytica*; and 1 ml/l of vitamin solution (Verduyn et al. 1992) and 10 ml/l of trace element solution (Meesters et al. 1996) for *R. toruloides*. For the auxotrophic strain *Y. lipolytica* Po1g, the MDBU medium was supplemented with 0.1 g/l of uracil and 0.1 g/l of leucine. The liquid medium MCL used for evaluating lipid accumulation capacity consisted of (g/l): glycerol, 70; yeast extract (containing 11.9% total nitrogen according to the manufacturer), 2.0; KH_2_PO_4_, 3.6; MgSO_4_.7H_2_O, 1.5; (NH_4_)_2_SO_4_,0.4; and trace element solution (Meesters et al. 1996), 10 ml/l. Total C/N molar ratio in the MCL medium was approximately 100 mol/mol. Both media were adjusted to pH 6 with KOH 1 mol/l.

### Microbial growth kinetics

The pre-inoculum was prepared from pouring a 1 ml stock culture (kept in glycerol at –80 °C) into a 500 ml baffled shake-flask containing 100 ml of YPD medium (10 g/l yeast extract, 20 g/l peptone, and 20 g/l glucose). The inoculum was prepared from 1 ml of the pre-inoculum in 100 ml of MDBU medium supplemented with 20 g/l of glycerol, thiamine solution for *Y. lipolytica*, and vitamin solution (Verduyn et al. 1992) for *R. toruloides*. Pre-inoculum and inoculum were cultivated in an orbital shaker at 200 rpm, 30 °C and 12 h for *Y. lipolytica* and 24 h and *R. toruloides* strains.

Cells from the inoculum were centrifuged at 2500 *g* for 10 min, washed twice with MDBU medium (without carbon source) and the cell suspension was standardized to start the growth kinetics experiment with an initial OD_600_ between 0.1 and 0.2 for *Y. lipolytica*, and 0.5 and 0.6 for *R. toruloides*. Cultivations were carried out in 500 ml baffled shake-flasks containing 100 ml of MDBU medium with 2.5 g/l of glycerol, thiamine and trace element solutions (Barth and Kunkel 1979) for *Y. lipolytica*and vitamin (Verduyn et al. 1992) and trace element (Meesters et al. 1996) solutions for *R. toruloides*, at 200 rpm and 30 °C.

### Cultivations for lipid accumulation

The pre-inoculum was prepared by pouring a 1 ml stock culture (kept in glycerol at –80 °C) into a 50 ml Falcon tube containing 10 ml of YPD medium and cultivation at 200 rpm and 28 °C for 24 h. The inoculum was prepared by transfering the 10 ml of pre-inoculum into a 500 ml baffled shake-flask containing 90 ml of YPD medium, and cultivation at 200 rpm and 28 °C for 24 h. Cells from the inoculum were centrifuged at 2500 *g* for 10 min, washed twice with NaCl solution (9 g/l) and the cell suspension was standardized to obtain an initial OD_600_ of 2.0 in the subsequent cultivation. Cultivations were performed in 500 ml baffled shake-flasks containing 100 ml of MCL medium at 200 rpm, 28 °C, and 96 h for *Y. lipolytica* and 72 h for *R. toruloides*. After this time, 10 ml of the broth were centrifuged at 2500 *g* for 10 min, washed twice with distilled water and the harvested biomass was dried at 100 °C until constant weight, which was later used for lipid quantification.

### Analytical Methods

Cell growth was monitored by optical density at 600 nm and converted into dry cell weight using a standard curve for each yeast strain. The dry weight of cells was determined gravimetrically, as described by Olsson and Nielsen (1997). Samples of the cultivations were filtered and the supernatants were used to quantify glycerol and extracellular metabolite concentrations using an HPLC equipment with an RI detector (RI 2000, Chrom Tech Inc., Germany) at 30 °C, Aminex HPX-87H ion exchange column (BioRad, USA) and H_2_SO_4_ 5 mM as mobile phase. Lipids were extracted from the dried biomass, obtained from the lipid accumulation experiments, according to Folch et al. (1957) and Bonturi et al. (2015).

### Calculation of physiological parameters

The maximum specific growth rate (μ_max_, h^−1^) was obtained by plotting the natural logarithm of dry weight (g/l) versus time (h), using only data points from the linear region corresponding to the exponential growth phase. μ_*max*_corresponded to the slope of the straight line obtained by linear regression. The biomass yield on substrate during the exponential growth phase (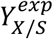, g DW/g S) was obtained by plotting the cell concentration (g DW/l) versus the glycerol concentration (g/l), using only data points from the exponential growth phase. 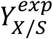corresponded to the absolute value of the slope of the straight line obtained by linear regression. The maximum specific substrate consumption rate (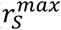), the global biomass yield on substrate (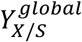), the lipid yield on substrate (*Y*_*Lip/S*_, g Lip/g S) and the lipid productivity (*P*_*Lip*_, g Lip/l.h) were calculated using the following equations:

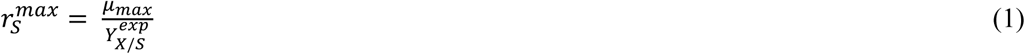

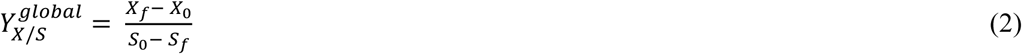

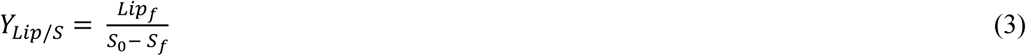

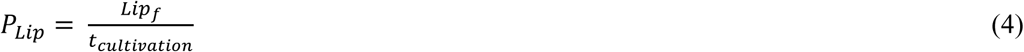

where *X*_*f*_ (g DW/l) is the final cell concentration, *X*_*0*_ (g DW/l) is the initial cell concentration, *S*_*f*_ (g/l) is the final glycerol concentration, S_*0*_ (g/l) is the initial glycerol concentration, *Lip*_*f*_ (g/l) is the final lipid concentration and t_cultivation_ (h) is the cultivation time.

### Statistical Analysis

The *Statistica® 5.5* (Statsoft, USA) software was used to calculate the analysis of variance (ANOVA). The Tukey test was used to determine the differences between the samples with a significance level of 10%.

## Results and discussion

The defined medium proposed by Barth and Kunkel (1979) and further modified by Oliveira (2014), herein called MDBU, was selected for the growth kinetics experiments with *Y. lipolytica*, due to its ability to: keep the extracellular pH constant, avoid foam formation, and sustain growth with a reasonably high specific growth rate (around 0.3 h^−1^) (Oliveira 2014). The growth profiles of all *Y. lipolytica* strains were in accordance with the typical microbial growth behavior in batch mode (Fig. 1) (Monod 1949) and as observed by Oliveira (2014). As for the *R. toruloides* CCT 0783 and

**Fig. 1.**
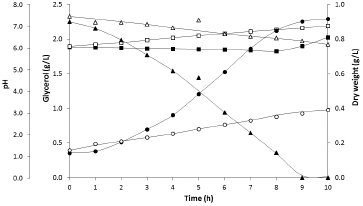
Cell concentration, glycerol concentration and pH along time of *Yarrowia lipolytica* IMUFRJ 50682 and *Rhodosporidium toruloides* CCT 7815 using glycerol as the sole carbon and energy source. Dry weight IMUFRJ 50682 (g/l - •), glycerol concentration IMUFRJ 50682 (g/l - ▲), pH IMUFRJ 50682 (▪), Dry weight CCT 7815 (g/l - ○), glycerol concentration CCT 7815 (g/l - Δ), pH CCT 7815 (□). The data points represent experimental data and lines represent trend lines. Results correspond to a typical replicate of three independent experiments.

*R. toruloides* CCT 7815 strains, the µ_*máx*_observed - 0,073 and 0,097 h^−1^, respectively – for cultivation in the MDBU medium proposed by Oliveira (2014) was roughly 3 times lower than the average μ_*máx*_value of the *Y. lipolytica* strains. For this reason, the MDBU medium was supplemented with a trace element solution (Meesters et al. 1996) and a vitamin solution (Verduyn et al. 1992), aiming at improvements in the µ_*máx*_of the *R. toruloides* strains. However, these modifications in the MDBU medium were not sufficient to increase this parameter, since the calculated µ_*max*_(with the modified MDBU medium) for *R. toruloides* CCT 0783 and *R. toruloides* CCT 7815 (Fig. 1) were 0.076 and 0.092 h^−1^, respectively.

According to Monod (1949), the growth of a microorganism can be more accurately measured when the limiting growth factors are fully known. These limiting growth factors can be classified into three groups: 1) depletion of nutrients; 2) accumulation of toxic metabolic products; or 3) change in the ionic balance, especially the pH. Thus, the use of a defined culture medium is important in the study of microbial physiology, as it is the only choice for acknowledging the limiting growth factors. In the case of *Y. lipolytica*, it was interesting to observe that the pH of the medium did not change along the cultivations, because urea was used as the nitrogen source and there was no production of organic acids during cell growth. Moreover, it was possible to conclude that glycerol was the limiting growth factor for this species, because the cells entered the stationary growth phase with the exhaustion of glycerol. However, this was not observed in the cultivations of *R. toruloides*: the extracellular pH increased during cell growth from 6.0 to 7.3 (for *R. toruloides* CCT 0783) or 7.0 (for *R. toruloides* CCT 7815), and the cell concentration seemed to stabilize before glycerol exhaustion. Therefore, it is possible that the increase in the extracellular pH was the limiting growth factor for the *R. toruloides* strains; however no explanations for this phenomenon or similar reports in the literature could be found.

Calculated physiological parameters and the kinects patterns observed during *Y. lipolytica* and *R. toruloides* cultivations in the defined medium are displayed in Table 1 and Fig. 1, respectively. Statistical analysis show that *Y. lipolytica* IMUFRJ 50678, *Y. lipolytica* Po1g, *Y. lipolytica* W29 and *Y. lipolytica* CCT 5443 can be considered physiologically very similar, because no significant difference (p < 0.1) was found, among these strains, for all the physiological parameters calculated (μ_*max*_ 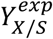 and 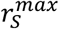). *Y. lipolytica* IMUFRJ 50682 presented no significant difference of µ_*max*_just with *Y. lipolytica* W29 strain, and in terms of 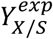and 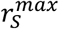a significant variation was found just with *Y. lipolytica* Po1g. As for the *Y. lipolytica* CCT 7401, significant difference was found for 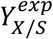with *Y. lipolytica* IMUFRJ 50678 and *Y. lipolytica* Po1g strains, and the µ_*max*_observed was significantly different from all the *Y. lipolytica* strains, but significantly equal in terms of 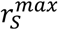.

The values presented in this work are in accordance with the findings of Santos et al. (2013) and Workman et al. (2013) for µ_*max*_- 0.18 h^−1^ and 0.32 h^−1^, respectively- and with Oliveira (2014) for 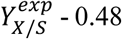 and 0.51 (g DW/ g S) - for *Y. lipolytica* strains cultivated in the defined medium using glycerol as a sole carbon and energy source. Oliveira (2014) also reported µ_*max*_and 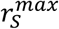values greater than what we observed in this work: 0.46 and 0.36 h^−1^ and 0.96 and 0.71 (g S / g DW.h) for *Y. lipolytica* IMUFRJ 50682 and *Y. lipolytica* W29, respectively.

Only *Y. lipolytica* W29 and both *R. toruloides* strains did not produce foam during cultivation for lipid accumulation. Foaming occurred during the first 24 h of culture when the nitrogen source was not yet completely consumed. Oliveira (2014) reported that excess of carbon and nitrogen sources in the medium were responsible for the foaming. Optical microscopy revealed that *Y. lipolytica* cells were present in the foam (data not shown), hindering the reproducibility of results, since those cells were not in contact with the liquid culture medium. The addition of antifoam in the cultivations can solve the excessive foam formation, but these additives may affect the growth of various microorganisms (Rosano and Ceccarelli 2014) and thus, should be avoided, whenever possible.

No significant difference (p < 0.1) was found for all analyzed physiological parameters when the two *R. toruloides* strains are compared. When compared to the *Y. lipolytica* strains, the µ_*max*_of *R. toruloides* CCT 0783 and *R. toruloides* CCT 7815 was, in average, 3.3 times lower, even after medium supplementation with vitamin solution (Verduyn et al. 1992) and trace element solution (Meesters et al. 1996). Most of the literature on *R. toruloides* aiming at lipids and/or carotenoids production reports the use of complex media to improve the maximum specific growth rate. Bommareddy et al. (2015) reported a maximum specific growth rate of 0.12 h^−1^ for *R. toruloides* DSMZ 4444 during a bioreactor cultivation using yeast extract and glycerol (75 g/l). Kiran et al. (2013) reported a maximum specific growth rate similar to the values found in this work, 0.085 h^−1^, during a bioreactor cultivation using pure glycerol (100 g/l) as the carbon source and autolysate of the rapeseed meal as nitrogen source.

Lipid accumulation capacity was evaluated using a complex medium with glycerol and a carbon to nitrogen ratio equal to 100 (mol/mol). Initially, the lipid accumulation kinectic was determined using *Y. lipolytica* IMUFRJ 50682 and *R. toruloides* CCT 7815 strains (Fig. 2). The lipid content, yield and productivity were subsequently determined for 72 h and 96 h of cultivation (highest lipid content) for the remaining *R. toruloides* and *Y. lipolytica* strains, respectively. *R. toruloides* strains were superior than *Y. lipolytica* strains in terms of *Y*_*Lip/S*_and *P*_*Lip*_(Table 1.). *Y. lipolytica* IMUFRJ 50682 showed the highest lipid content (59.29%), but presented the lowest *Y*_*Lip/S*_and *P*_*Lip*_values along with a low biomass concentration (average of 1.3 g DW/l), therefore discouraging its application in industrial lipid production processes.

**Fig. 2.**
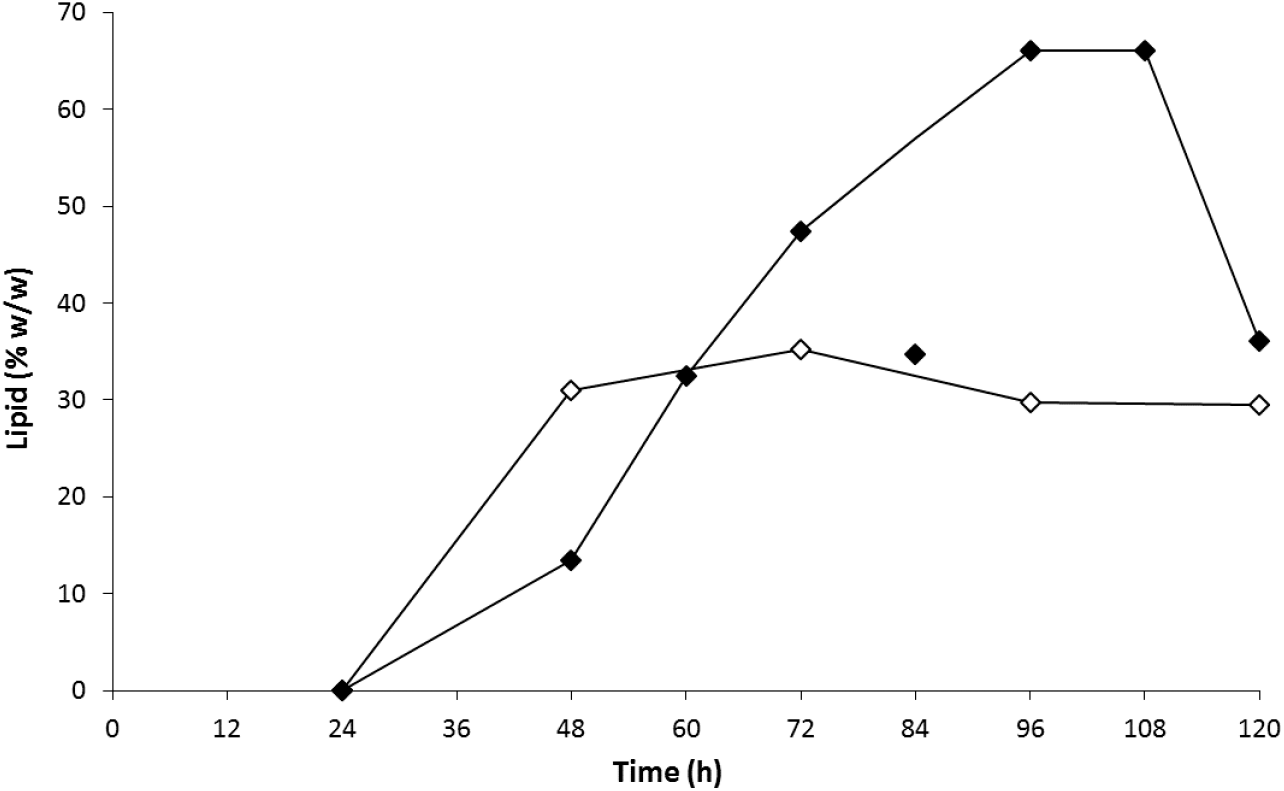
Kinetics of lipid accumulation in a complex medium of *Yarrowia lipolytica* IMUFRJ 50682 and *Rhodosporidium toruloides* CCT 7815 using glycerol as the sole carbon and energy source. Lipid percentage IMUFRJ 50682 (% - ♦) and lipid percentage CCT 7815 (% - ◊). The data points represent experimental data and lines represent trend lines. Cultivation was performed without replicates.

*R. toruloides* CCT 7815 is an evolved strain of the wild-type *R. toruloides* CCT 0783. The evolution was carried out using successive cultivations in increasing concentrations of hydrolysed sugarcane bagasse (Bonturi et al. 2017). In this study, Bonturi et al. (2017) cultivated both *R. toruloides* strains in complex medium with glucose, xylose or a mixture of xylose:glucose at a mass ratio of 6:1 as the carbon source, and reported the tendency of the *R. toruloides* adapted (CCT 7815) to accumulate more lipid than the parental strain (CCT 0783) in the three carbon sources (Table 1).

Physiological characteristics of the evolved strain, in this work, were not statistically different from the ones displayed by the parental strain (Table 1); however, *R. toruloides* CCT 7815 strain was superior not only to the parental strain, but to all other strains studied here, in terms of lipid production. These results also suggest that the evolution process to which the *R. toruloides* CCT 0783 strain was submitted did not change the basic physiological characteristics of this strain, but modified lipid accumulation characteristics, possibly as a response to the increasing stress applied along the evolution strategy. The evolution process was performed at 28 °C and 200 rpm, in medium containing 20.0 g/l xylose, 5.0 g/l glucose, 1.9 g/l yeast extract, 1.5 g/l MgSO_4_·7H_2_O, 5.0 g/l (NH_4_)_2_SO_4_, 3.6 g/l KH_2_PO_4_, and 2% trace minerals solution. After reaching the exponential growth phase, 25 ml of broth were centrifuged and cells were resuspended in fresh medium with addition of 10 % (v/v) of the hydrolysed sugarcane bagasse. For each step of the evolution process, the proportion of hydrolysate in the medium was increased by 10 % until the yeast was capable to grow in 100 % of hydrolysate (Bonturi et al 2017). As for the parental *Y. lipolytica* W29 and the modified *Y. lipolytica* Po1g strains, both physiological and lipid accumulation characteristics did not change despite the genetic manipulations present in Po1g.

Stoichiometric calculations showed that the theoretical maximum conversion factor of substrate into lipids produced by oleaginous microorganisms is 0.32 g Lip/g S, 0.34 g Lip/g S, and 0.30 g Lip/g S for the substrates glucose, xylose and glycerol, respectively. However, experimental *Y*_*Lip/S*_values around 0.10 g Lip/g S are typically reported (Papanikolaou and Aggelis 2011). The maximum value reported was 0.27 g Lip/g S (Bommareddy et al. 2015), on pure glycerol (75 g/l) as carbon source.

## Conclusions

From our data, albeit presenting three times lower specific growth rates than *Y. lipolytica*, the yeast *R. toruloides*, specially the strain *R. toruloides* CCT 7815, presents greater potential to be used for industrial SCO production than the former yeast, due to its ability to convert glycerol into lipids close to the experimental *Y*_*Lip/S*_values typically reported for oleaginous microorganisms (0.11 g Lip/g S) and for having displayed the highest lipid productivity among all strains studied here, 0.10 (g Lip /l.h). Furthermore, this species is also able to produce carotenoids, a high value product, concomitantly to SCO, which could be explored to facilitate the achievement of economical viability in microbial oil production processes (Koutinas et al. 2014).

## Acknowledgements

This work was financed by Conselho Nacional de Desenvolvimento Científico e Tecnológico (CNPq, Brasília, Brazil), process number 130496/2014-6. Everson Alves Miranda and Nemailla Bonturi thank the Fundação de Amparo à Pesquisa do Estado de São Paulo (FAPESP, São Paulo, Brazil), grant number 2013/03103-5, for financial support. We would like to thank Prof. Maria Alice Zarur Coelho, Federal University of Rio de Janeiro, for providing *Yarrowia lipolytica* strains.

